# Adaptive selection of p53 mutation metaplastic phenotypes in estrogen-independent progression of ER+ tumors: A mechanism for acquired resistance to hormonal therapy

**DOI:** 10.1101/2025.02.02.636128

**Authors:** Hyeongsun Moon, Hidetoshi Mori, Jane Chen, Alejandra Patino, Zsofia Penzvalto, Kavya Ramamurthy, Jaeyong Choi, John D. McPherson, Joshua C. Snyder, Robert D. Cardiff, Alexander D. Borowsky

## Abstract

Estrogen receptor positive (ER^+^) subtypes of mammary adenocarcinoma comprise 79% of all breast cancer diagnosis and 67% of all breast cancer mortality. The paucity of models of ER^+^ mammary cancer that mimic human disease and response to treatment has limited critical preclinical study of mechanisms and new therapies for ER^+^ breast cancer. The *Stat1* knockout, 129S6/SvEvTac-Stat1^tm1Rds^ (*Stat1^-/-^*), females develop luminal type FoxA1^+^, ER^+^, and PR^+^ mammary carcinomas after prolonged latencies. Initial studies showed that a cell line derived from a Stat1-/- mammary carcinoma was tumorigenic in syngeneic mice, but non-tumorigenic in ovariectomized (Ovx) mice. Here, data shows that Ovx performed after SSM2 tumors establish growth results in ovarian hormone independent growth. The viable post-Ovx tumors were primarily composed of metaplastic CK14^+^ basal type cells with a high percentage p53 immunohistochemistry (IHC) positive “mutation pattern”, rather than the original luminal type tumors with low percent “wild type” pattern p53. Comparing whole exome sequences of ER^+^ *Stat1^-/-^* mammary tumors before and after Ovx, revealed basal keratins, mesenchymal (EMT) phenotypes, and unique mutation profiles in genes, including *Trp53* and *Prlr,* in the estrogen-independent tumors. Our experimental findings are consistent with the clinical evidence of tumor heterogeneity of ER^+^ breast cancers in patients in recent whole genome sequencing studies. Similarly, spontaneous Stat1-/- tumors with high percentage p53 “mutation pattern” were more basaloid and grew rapidly after Ovx, while retaining high expression of ER and FoxA1. This study demonstrates that the STAT1^-/-^, ER^+^ estrogen dependent breast cancers can become resistant to through clonal selection of mammary cells comprised of metaplastic p53^+^/CK14^+^ basaloid cells.

## INTRODUCTION

The genomic era has led to major advances in the diagnosis, treatment, and cure of cancers. However, therapeutic resistance and recurrence in many cancers, including estrogen-receptor-positive (ER^+^) breast cancer, remain problematic (1–3). Studies using genetically modified mouse models (GEMM) have provided various hypotheses including oncogene addiction (4–7), adaptive oncogenesis (8, 9), biological predeterminism (10, 11) and metaplastic reprogramming (12–14) as potential mechanisms of development of drug-resistant cancers. The development of mouse model systems has also encouraged a search for reproducible ER^+^ mammary cancers, although the lack of robust ER^+^ mammary tumor models in GEMM has been a limiting factor (15–19).

129S6/SvEvTac-Stat1tm1Rds (*Stat1^-/-^*) mice develop ER^+^ mammary tumors which resemble luminal type breast cancer in human patients (20). Mammary tumors in *Stat1^-/-^* mice arise late in age, average 18 months and are incompletely penetrant in this model (20, 21). Tumors in this model have been previously reported to arise after spontaneous heterozygous Prlr truncating mutations (22), and tumor growth is reported to be estrogen-dependent (20, 23). A series of cell culture lines derived from these tumors were developed (20), and one of these, the “SSM2” cells have been characterized in our laboratory. They are tumorigenic when injected into the mammary fat pad of syngeneic, immune intact 129S6/SvEv mice, develop ER+ FOXA1+ adenocarcinomas, and express both immune-attractive and immune-repulsive factors resulting in an “immune-excluded” phenotype(21). Similar to these SSM2 cell lines, fresh biopsies derived from the spontaneous *Stat1^-/-^* mammary tumors are tumorigenic in orthotopic syngeneic transplantation

Although previous studies show that ovarian estrogen is necessary for SSM2 cell tumorigenesis in pre-ovariectomized mice (20), we reasoned that a more important test of hormone-sensitivity would require an experimental design more representative of the human clinical condition under hormone therapy. Specifically, we asked whether the ablation of ovarian estrogen *after* tumors had developed and were measurable, would control or reverse the growth and/or progression of established tumors (Supplementary figure 1).

We describe herein that ovariectomy following transplantation and established growth of *Stat1*KO mouse derived primary mammary tumors and mammary tumor cell line resulted in continued tumor size. However, histological examination revealed the loss of the luminal cell tumor population in post-ovariectomy tumors and the emergence of metaplastic, basaloid tumor cell populations. DNA Sequence analysis proved that the post-ovariectomy metaplastic tumors were composed of cells with unique mutation profiles in both the *Trp53* and *Prlr* genes. In a second set of experiments, we instead utilized transplanted biopsies of spontaneous tumors in the *Stat1*-/- mice and intentionally included one common tumor type with low percent p53 by IHC and a second uncommon primary tumor with high percent p53 for comparison.

## RESULTS

### *Established SSM2* cell tumors transplants adapt to loss of ovarian estrogen

SSM2 cells establish tumors in immune and ovary intact syngeneic 129S6SvEv mice and show ER and PR expression with very low percent p53 (wild-type pattern) by IHC (Figure 1a). In this experiment, SSM2 cells were transplanted into the mammary fat pads of intact female syngeneic 4-week-old 129S6SvEv mice (Figure 1b and Supplementary Fig. 1). At day 29 (4 weeks post-transplantation), all tumors were palpable and measurable with calipers, though none exceeded 3 mm in largest diameter. Ovariectomy was performed on day 29, and tumor size was monitored over the following 8 weeks. Tumors were harvested at day 44 (2 weeks post-ovariectomy), day 65 (5 weeks), and day 86 (8 weeks; endpoint) for histological analysis, immunohistochemistry, and gene sequencing. At harvest, two distinct phenotypes were observed. Although both tumors were ER+, one phenotype exhibited a basaloid appearance, increased basal keratin (CK14) expression, and a high percentage of p53-positive cells (Figure 1c,d). In contrast, the other phenotype displayed a luminal appearance with a low CK14 expression and a low percentage of p53-positive cells. Growth curves were plotted based on phenotype (Figure 1e), revealing distinct differences in post-ovariectomy growth rates (Figure 1f). Tumors with a luminal phenotype established growth more rapidly but plateaued significantly after ovariectomy. Conversely, tumors with a basaloid phenotype exhibited slower initial growth but maintained or even accelerated growth following ovariectomy (Figure 1e).

**Figure 1.**
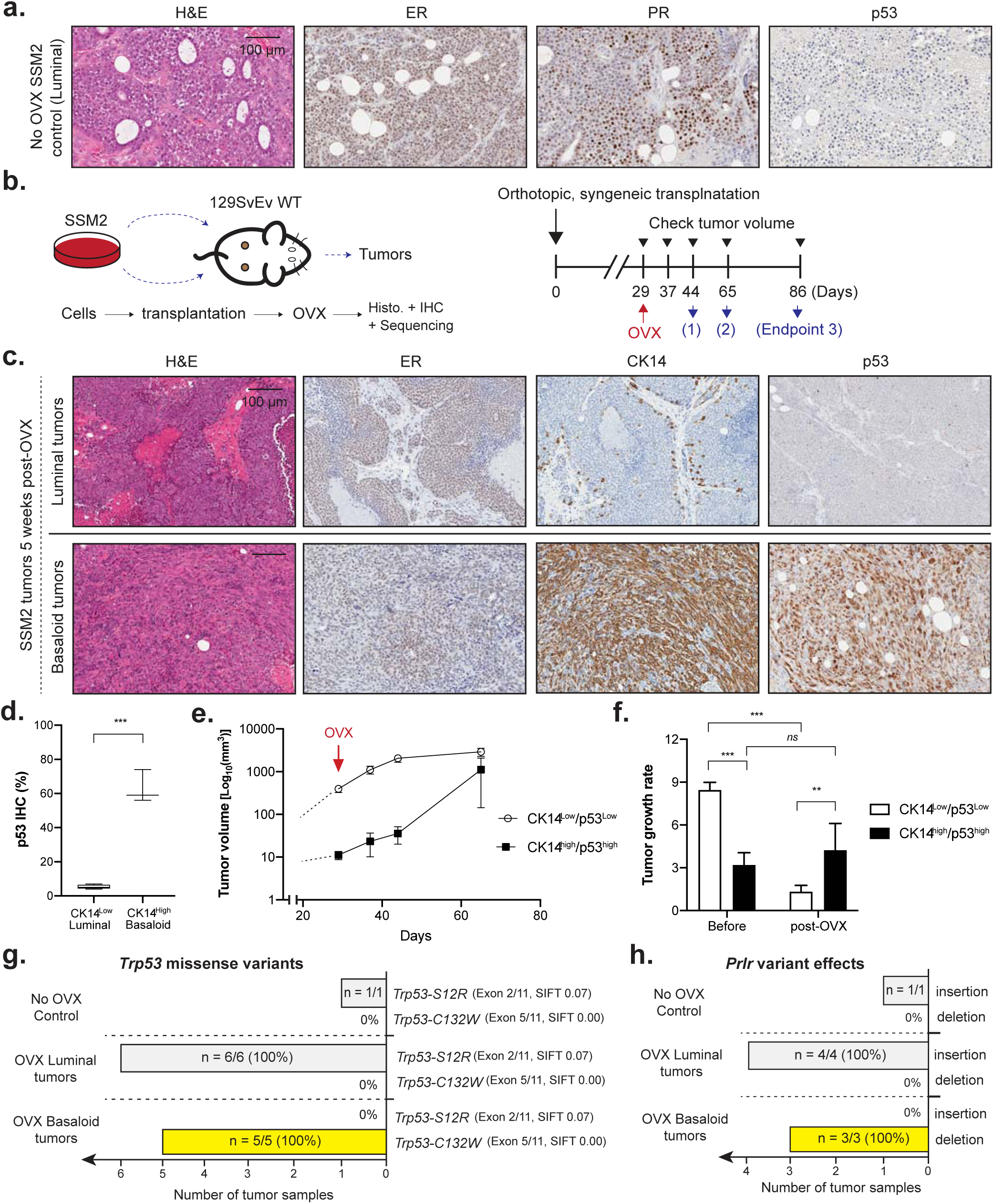
Depletion of estrogen triggered a phenotypical conversion from luminal tumors to p53-high basaloid tumors in syngeneically orthotopically transplanted luminal-type *Stat1*-null SSM2 cells. (**a**) The *Stat1*-knockout mouse-derived cell line, SSM2, formed hormone receptor-positive mammary tumors when syngeneically orthotopically transplanted into wild-type mice. Shown images are tissue sections stained with H&E, estrogen receptor (ER), progesterone receptor (PR) and p53. (**b**) A schematic representation of the ovariectomy (OVX) procedure and tumor size monitoring. Tissue isolations were performed at 2 weeks (1), 5 weeks (2), and 8 weeks (Endpoint 3). (**c**) Phenotypic conversion from luminal tumors to basaloid tumors was observed in SSM2 tumor transplants at 5 weeks post-ovariectomy. Shown images include H&E, ER, cytokeratin 14 (CK14), and p53 for luminal tumors (upper panel) and basaloid tumors (lower panel). (**d**) Immunohistochemical analysis for p53 positive tumor cells shows an increase in p53-positive tumor cells in basaloid (CK14-high) tumors. (**e**) Tumor volume measurements after OVX treatment depict growth curves for CK14-low/p53-low luminal tumors (white squares) and CK14-high/p53-high basaloid tumors (black squares). (**f**) Tumor growth comparisons of CK14-low/p53-low and CK14-high/p53-high tumors before and after OVX treatment. (**g**) Gene mutation analysis revealed that the p53 (*Trp53*) missense mutation C132W dominated the cell population in the basaloid tumor after ovariectomy treatment. (**h**) Variant mutations in the prolactin receptor also shifted from luminal tumors to basaloid tumors following ovariectomy treatment. Scale bar=100 microns. Asterisks indicate statistical significance: *** (p<0.001) and ** (p<0.01) based on t-test.

Gene sequencing analysis of these tumors confirmed the surrogate IHC assay results, showing that a high percentage of p53-positive cells by IHC correlated precisely with a *Trp53* p.C132W mutation. This mutation is analogous to a known deleterious hotspot mutation in human tumors. Meanwhile the low percentage p53 tumors were found to have a *Trp53* p.S12R variant, which is predicted by SIFT score (24) to be non-functional (Figure 1g). As expected, there was also a truncating alteration in *Prlr* in both tumor types, but the luminal tumor alteration was a specific insertion, whereas the basaloid tumor alteration was a deletion. Identical *Trp53* variants and *Prlr* variants were seen in all the basaloid tumors, suggesting an origin from a subclone of the parent SSM2 cells transplanted rather than spontaneous mutations arising after ovariectomy. Additional confirmation of coexistence of these two subclones in the SSM2 cell line, with adaptive segregation and preferential growth after ovariectomy is provided by the whole exome sequence summarized (Figure 4) and discussed below.

### Estrogen-independent growth of SSM2 tumors emerges with CK14+/p53high clones

To study the progression of SSM2 tumors toward estrogen-independent growth, we compared tumors harvested early time point (2 weeks post-ovariectomy) with those collected at the 8-week endpoint. This comparison revealed the emergence of CK14+ foci at 2 weeks (Figure 2a), which expanded to occupy the entire tumor area by 8 weeks (Figure 2b). Multiplex staining was performed to simultaneously assess ER, CK14, p53, and the proliferation marker Ki67 in tumor tissues from both the 2-week (Figure 2c) and 8- week (Figure 2d) time points. This approach allowed us to evaluate the proliferative rates of distinct cell subsets within the tumors. The proportion of CK14+ cells increased significantly, rising from 25% with mid-level expression at 2 weeks to 83% with mid-level and an additional 7% with high-level expression at 8 weeks (Figure 2c,d). Segregating Ki67 expression by cell type, based on CK14 levels, revealed that CK14+ cells are significantly more proliferative than CK14-negative cells. Moreover, cells with higher CK14 expression demonstrated the highest proliferative rate, with 52% Ki67 positivity (Figure 2c right, 2d right). Summary tables (Figures 2e and 2f) comparing the CK14 low and CK14 high areas’ cell proliferation indicate that cells with higher levels of CK14 were increased both luminal- and basaloid tumor but the basaloid tumor had more CK14 cell proliferation. The proliferation status of p53- and p53+ cells in 2wks post-OVX tumor (Figure 2g) and in 8wks post-OVX tumor (Figure 2f) showed that p53+ cells are more proliferative compared with p53-cells in both conditions. Notably, ER is still expressed in this CK14+/p53+ proliferative cell type at 2 weeks (Figure 2i), but a subset of these cells at 8 weeks may be emerging with 15% negative for ER (Figure 2j). Overall, this supports the model that the ER+/CK14-/wild type p53 cell growth is favored in the presence of estrogen, but the ER+/CK14+/mutant-p53 cell growth is favored after ovariectomy (Figure 2k).

**Figure 2.**
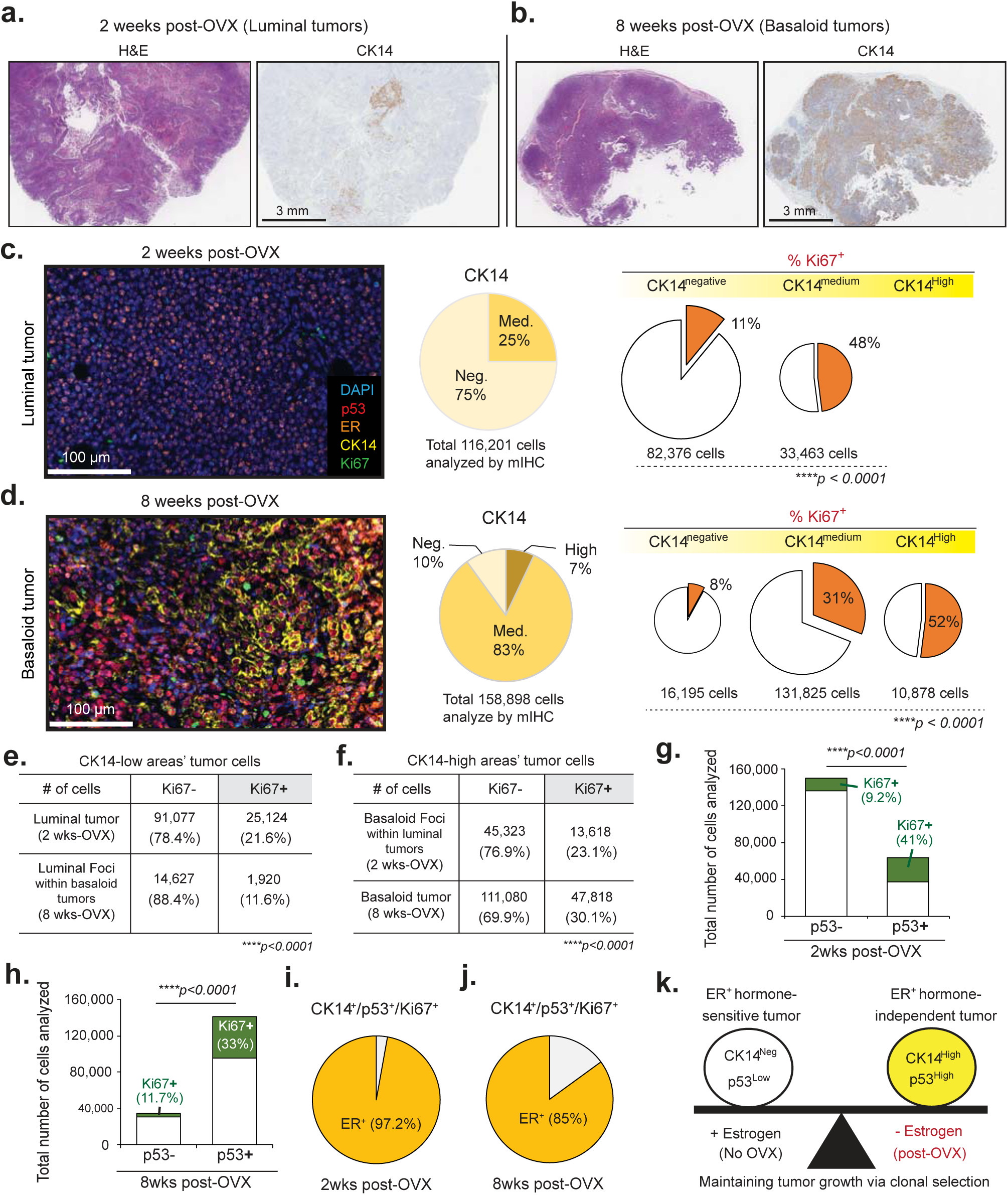
SSM2 tumors undergo phenotypic conversion from luminal to basaloid tumors following ovariectomy (OVX) treatment. SSM2 tumor transplants at two and eight weeks post-OVX were compared using immunohistochemical analysis. (**a, b**) SSM2 tumors transitioned from luminal to basaloid phenotypes post-OVX, as shown in H&E and CK14 IHC staining for tumors at (**a**) two weeks and (**b**) eight weeks post-OVX. (**c, d**) Multiplex immunohistochemical analysis for p53 (red), ER (orange), CK14 (yellow), Ki67 (green), and DAPI (blue) was performed to examine proliferating cell types in (c) luminal tumors (two weeks post-OVX) and (**d**) basaloid tumors (eight weeks post-OVX). CK14-positive cells were analyzed by expression level (negative, low, medium, high) and proliferation status, revealing that basaloid (CK14-positive) cells proliferate more than CK14-negative cells at both time points. (**e, f**) Comparisons of Ki67+ tumor cells at CK14-low (**e**) and CK14-high (**f**) regions indicate higher proliferation in luminal tumors at two weeks post-OVX and in basaloid tumors at eight weeks post-OVX. (**g, h**) Proliferation of p53-positive and p53-negative cells was compared, showing higher proliferation in p53-positive cells under both two-week (**g**) and eight-week (**h**) post-OVX conditions. (**i, j**) Pie charts illustrate the percentage of ER-positive cells within CK14+/p53+/Ki67+ populations at (**i**) two weeks and (**j**) eight weeks post-OVX. (**k**) A schematic representation summarizes the phenotypic transition of SSM2 tumors in the presence and absence of OVX treatment. Asterisks indicate statistical significance: *** (p<0.001) based on the Chi-square test.

### Spontaneous tumors in the *Stat1*-/- mouse sometimes arise with *Trp53* mutations and estrogen independence

The typical mammary carcinoma arising in the *Stat1*-/- mouse is a luminal phenotype adenocarcinoma with wild-type *Trp53*. However, we observed a minority of tumors with a more basaloid phenotype, and upon investigation these were found first to be high percentage p53 IHC positive and then found to harbor *Trp53* gene variants corresponding to several hot-spot mutations seen in human cancers. Both tumor types express ER and PR (Figure 3a). Since we routinely cryopreserve viable, transplantable biopsies of harvested tumors, we were able to conduct an additional ovariectomy experiment using biopsies from these two tumor types. The luminal/p53-low tumor was transplanted into the right mammary fat pad, while the basaloid/p53-high tumor was transplanted into the left (Figure 3b). Similar to the SSM2 transplant experiment, ovariectomy was performed after the tumors had established growth and were measurable with calipers at 5 weeks post-transplant. Tumors were allowed to grow for another 2.5 weeks and then harvested for analysis (Figure 3b). The comparative tumor growth before and after ovariectomy is shown (Figures 3c-d). The growth rate of the basaloid *Trp53*-mut tumor increases significantly after ovariectomy, whereas as the lumina tumor plateaus completely with a near zero growth rate after ovariectomy (Figure 3c). Post ovariectomy phenotypes of ER and PR were retained at a similar level (Figure 3f). Additional multiplex IHC analysis for visualizing CK14, ER, p53 and Ki67 (Sup. Figure 3a,b) shows that the p53high tumor has overall a higher percentage of CK14+ cells and the cell proliferation is higher in CK14+ cells in both p53high and p53low tumors (Sup Figure 3b). Interestingly, the proliferative rate did not obviously differ between p53+ and p53− cells in either tumor type, nor between the p53-high and p53-low tumors (Sup. Figure 3c), suggesting that the residual viability in the luminal p53-wt was not a product of emergence of a new spontaneous p53 mutation leading to a new basaloid proliferative phenotype. It is possible that such a spontaneous subclone might emerge if given a longer time. Note also, that both the luminal and basaloid primary tumors express almost equal percentages of ER+ cells after ovariectomy (Sup. Figure 3d).

**Figure 3.**
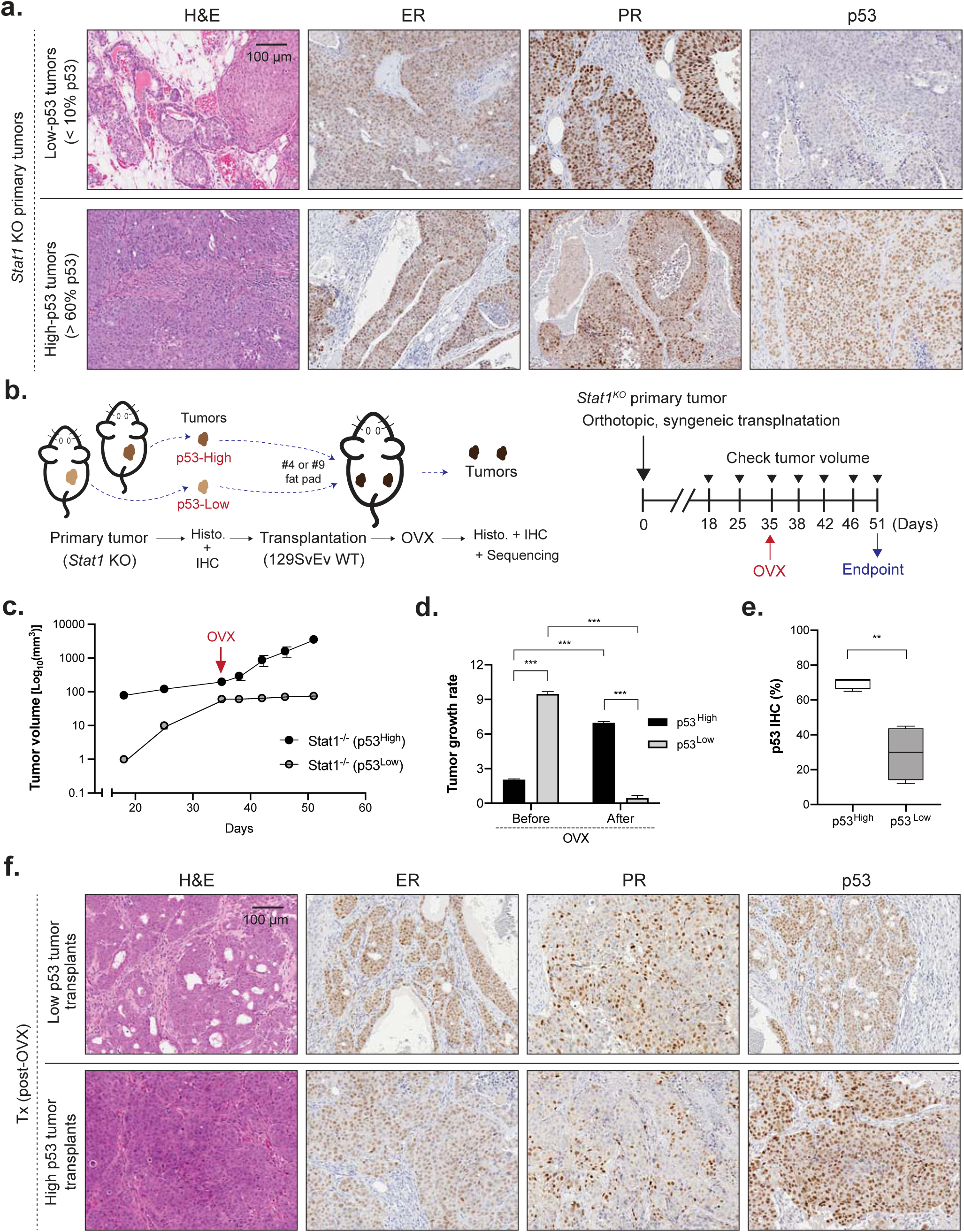
*Stat1* knockout primary tumor transplants exhibit differential growth based on ovarian hormone status. (**a**) *Stat1*-null mammary tumors in *Stat1* knockout mice display distinct differences in p53 status. While these tumors are predominantly ER+/PR+, p53 expression varies, as shown in representative images of low-p53 (upper panel) and high-p53 (lower panel) tumors. (**b**) A schematic representation outlines the experimental procedure for comparing the growth of p53-high and p53-low *Stat1*-null primary mammary tumor transplants before and after ovariectomy (OVX). *Stat1*-null primary tumors were isolated, their p53 status was assessed, and the tissues were syngeneically orthotopically transplanted into wild-type mice. OVX treatment was performed at day 35, and tumors were collected 51 days post-transplantation. (**c**) Tumor growth curves for p53-high (black) and p53-low (white) *Stat1*-null tumor transplants are shown. (**d**) Comparative analysis of tumor growth rates before and after OVX treatment between p53-high and p53-low *Stat1*-null tumor transplants. (**e**) Immunohistochemical analysis of p53 expression in p53-high and p53-low tumor transplants at the experimental endpoint. (**f**) Histological and immunohistochemical comparisons of p53-low and p53-high Stat1-null tumor transplants after OVX treatment. Shown images include H&E staining and IHC for ER, PR, and p53 in p53-low tumors (upper panel) and p53-high tumors (lower panel). Asterisks indicate statistical significance: *** (p<0.001) and ** (p<0.01) based on t-test.

### The whole exome sequence reveals at least 2 distinct clones in SSM2 cells, demonstrates that additional *Trp53* hot-spot mutations confer estrogen independence, and reveal new spontaneous variants that may reflect alternative mechanisms of estrogen independence

Our initial analysis using whole exome sequencing revealed unique additional mutations in cancer driver genes between luminal and basaloid SSM2 tumors post-ovariectomy compared to control SSM2 tumors (Supplementary Table 1). Interestingly, from the whole exome or Sanger sequencing analyses, deleterious *Trp53* mutations were found in hormone-independent, metaplastic basaloid tumors (Table. 1). Specifically, SSM2 basaloid tumors express *Trp53* p.C132W which is considered to have an identical effect as a deleterious mutant *Tp53, Tp53 p.C135W,* in human patients (Table 1). *Tp53* p.C135W is common in human cancer patients including invasive breast ductal adenocarcinoma and high grade ovarian serous adenocarcinoma (25). Notably, non-deleterious *Trp53* mutations found in luminal tumors were not detected in any basaloid tumors (Table. 1). Second, mammary tumorigenesis can also be promoted by persistent PRLR signaling (23), and it has also been reported that *Prlr* truncation can also be associated with increased *in vivo* growth of ER^+^ mammary tumors in *Stat1^-/-^* mice (22). In addition to *Trp53* missense variants, basaloid tumors showed different *Prlr* mutations compared to luminal mammary tumors (Table. 2). Notably, the identical *Trp53* p.C132W and identical *Prlr* variants were seen in each phenotype, strongly suggesting that the *Trp53* p.C132W subclone is present, albeit at a low frequency, in the parent SSM2 cell line. It would be unlikely that both variants would co-occur spontaneously in all three samples. In subsequent fluorescent tagging experiments, we were able to separate these subclones (shown in Figure 5).

To investigate differences in gene mutations in SSM2 transplant tumors, we compared four experimental groups: OVX-treated SSM2 tumor transplants with low p53 expression (Group 1: p53low/ovariectomized) and high p53 expression (Group 2: p53high/ovariectomized), *Stat1*KO-derived mammary tumors without OVX treatment (Group 3: *Stat1*KO/intact), and SSM2 tumor transplants without OVX treatment (Group 4: SSM2/intact). Since the SSM2 cell line was the origin of this experiment, we hypothesized that tumors without OVX treatment (Group 4) represent parental gene mutations, while tumors under OVX treatment (Groups 1 and 2) may exhibit adapted cell populations or newly acquired mutations after OVX treatment. Gene mutations were categorized into two groups: those mutated (Figure 4.a) and those not mutated (Figure 4.b) in SSM2 tumors without OVX treatment (Group 4). The analysis revealed distinct mutation patterns between p53low and p53high SSM2 tumor transplants under OVX treatment. To summarize the similarities and differences, the mutated genes were organized into five clusters (Ca1–Ca5) which were positive for Group 4 (Figure 4a), and six clusters (Cb1–Cb6) which were negative for Group 4 (Figure 4b). These clusters highlight patterns of gene mutations across the experimental groups, facilitating the identification of mutations specific to OVX treatment, p53 expression levels, and the *Stat1*KO or SSM2 tumor background. Grouping genes based on their presence or absence in Group 4 (SSM2 tumors without OVX treatment) provided insights into parental gene mutations, OVX-dependent mutations, and those associated with specific tumor phenotypes or conditions.

**Figure 4.**
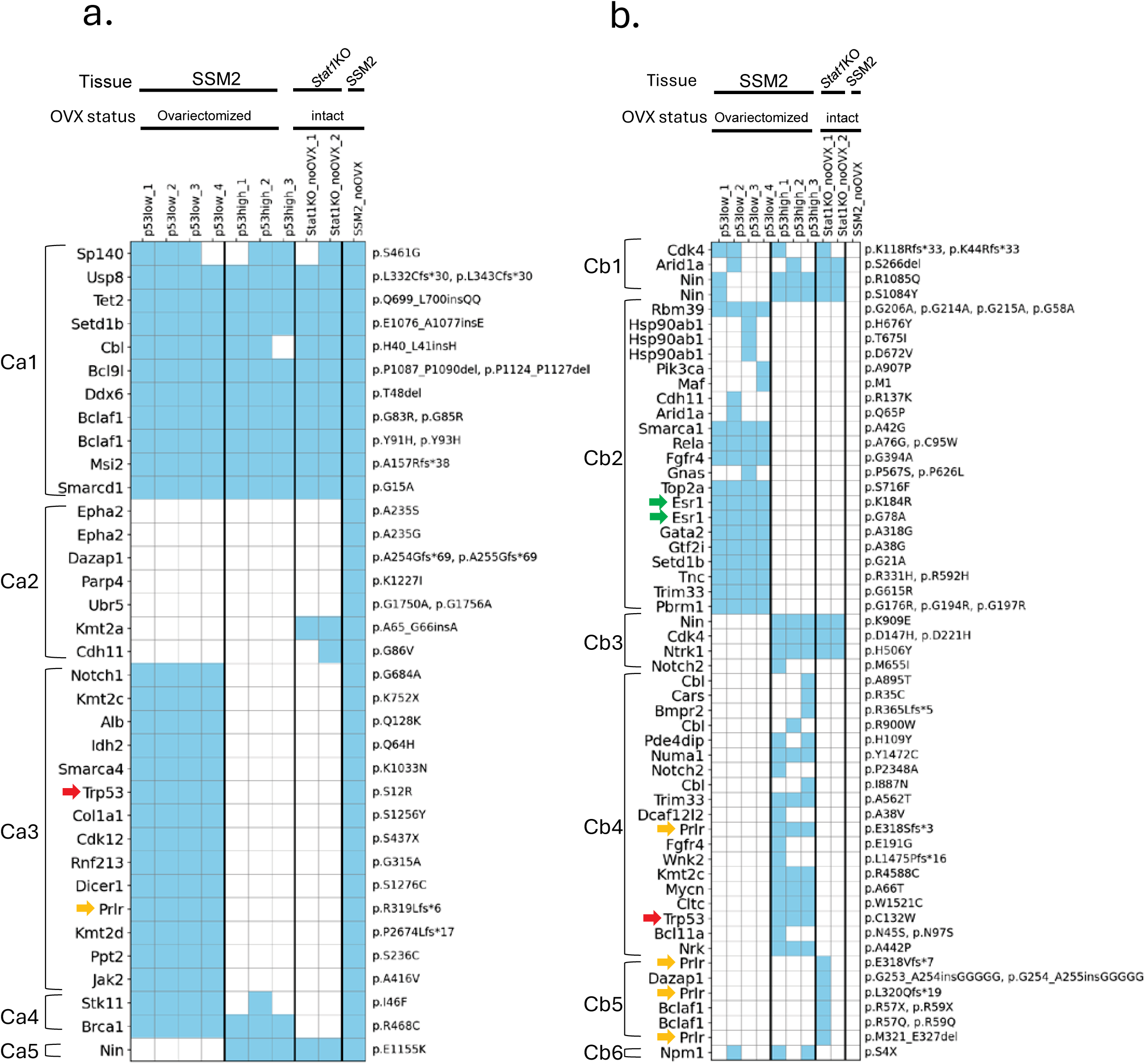
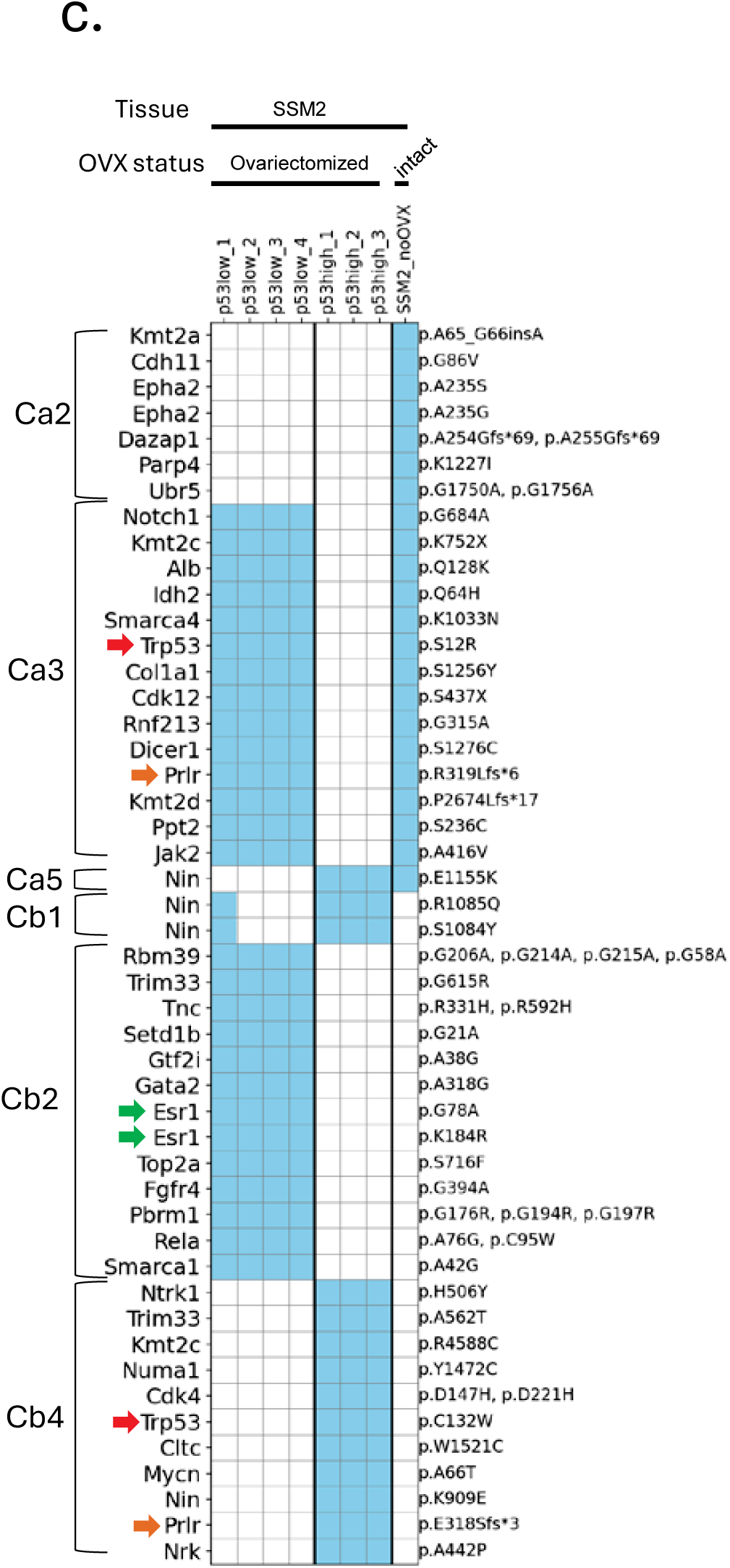
Gene mutation analysis in p53low and p53high SSM2 tumors under ovariectomized conditions. OVX-treated SSM2 tumors with low p53 expression (p53low1–4; Group 1) and high p53 expression (p53high1–3; Group 2) were analyzed for gene mutations. Tumors from *Stat1*KO mice without OVX treatment (Stat1KO_noOVX1&2; Group 3) and SSM2 tumors without OVX treatment (SSM2_noOVX; Group 4) served as reference samples and were included for comparison. The plots depict (a) genes positive for mutations and (b) genes negative for mutations in SSM2 tumors without OVX treatment. Blue squares indicate mutation-positive cases. Gene names are displayed on the left, with the corresponding mutation description on the right. Mutation clusters are grouped as Ca1–Ca5 in (a) and Cb1–Cb6 in (b). Arrows highlight key mutations: *Trp53* (red), *Prlr* (orange), and *Esr1* (green). (c) Unique patterns of gene mutations in each SSM2 tumor characterizing the cell population.

Mutations in the Ca1–Ca5 clusters were observed in all groups, indicating that these mutations are pre-existing in the SSM2 tumor transplant, present in the *Stat1*KO primary tumor, and persist in the SSM2 tumor under OVX conditions (Figure 4a). Mutations in Ca2 were consistently present in Group 4 and occasionally observed in Group 3, indicating these may be unique to *Stat1*KO mammary tumor without OVX treatment. In addition, mutations in Ca2 were not observed in OVX treated SSM2, suggesting this Ca2 might be specific to the condition in the presence of ovarian hormones. Genes in Ca3 were positive in Groups 1 and 4 but negative in Groups 2 and 3, implying that these mutations may link to SSM2 tumor transplants under p53low conditions. Ca4 mutations were observed in Groups 1, 2, and 4 but absent in Group 3, indicating that these are specific to SSM2 cells. One mutation in Ca5 was positive in Groups 2, 3, and 4 but not in Group 1, suggesting these are associated with either the p53high phenotype or basaloid characteristics.

In the Cb1–Cb6 clusters, mutations in these categories were absent in Group 4, suggesting that SSM2 tumor transplants generated these mutations under OVX condition (Figure 4b). Mutations in Cb1 were present in Groups 1 and 2, indicating they arise in SSM2 tumors after OVX treatment. Mutations in Cb2 were observed only in Group 1, indicating that they are specific to p53low SSM2 cells under OVX conditions. Interestingly, two mutations in the estrogen receptor-1 (*Esr1*) AF1 domain (p.G78A) and DBD domain (p.K184R) were found in this cluster, suggesting mutant Esr1 may play a role in estrogen-independent functions. Mutations in Cb3 and Cb4 were positive in Group 2 but negative in Group 1, suggesting that these genes may contribute to phenotypic conversion from luminal to basaloid cell types in SSM2 tumors under the OVX condition. Cb5 mutations were observed exclusively in *Stat1*KO tumors (Group 3) but only one line of *Stat1*KO tumor was positive for these mutations. One mutation in Cb6 was observed in Groups 1 and 2, indicating it may be involved in OVX treatment-dependent cellular processes. Additionally, a hybrid analysis of tumors Ca1–Ca5 and Cb1–Cb6 revealed how p53-high and p53-low tumors adapted cell populations with specific mutations (Ca2, Ca3, Ca5, Cb1, Cb2, Cb4), both with and without OVX treatment (Figure 4c). Consistent with the Trp53 and Prlr mutation patterns summarized in Figures 1g and 1h show that other genes in Ca3 and Cb4 exhibited similar trends, with Ca3 corresponding to luminal tumors and Cb4 to basaloid tumors. Additionally, the figure highlights distinct cell populations responding to ovariectomy: both p53-low and p53-high tumors (Ca2), p53-low luminal tumors (Cb2), and p53-high basaloid tumors (Cb4). Regarding Nin mutations, Ca5 and Ca1 display notable patterns. Ca5 mutations were present in both SSM2_noOVX and p53-high basaloid tumors, while mutations in Cb1 were associated with ovariectomy-responsive cell populations, predominantly observed in p53-high basaloid tumors.

These results from Figure 4 indicate that SSM2 tumors developed with different lineages of SSM2 cells carrying specific mutations or by generating new mutations under the OVX condition.

### SSM2 cells segregate into estrogen-dependent and estrogen-independent subclones

To investigate this, the SSM2 cell line was stably transfected with a series of fluorescent protein expression vectors driven by a ubiquitous CMV promoter (Figure 5a–c). Following transfection, serial flow sorting was used to isolate cells based on fluorescent reporter expression, yielding four distinct color-coded subclones of SSM2 cells (Figure 5d). These subclones were cultured in standard complete media (estrogen-replete) and estrogen-depleted (charcoal-stripped) media. After 8 days without estrogen, three of the color clones (EYFP, FR-MQV, and mTFP1) either died or underwent senescence. In contrast, one color clone (mKO) maintained growth and viability in estrogen-depleted conditions (Figure 5d), suggesting that this transfection targeted a cell from the low-frequency estrogen-independent subset of the mixed SSM2 culture.

**Figure 5.**
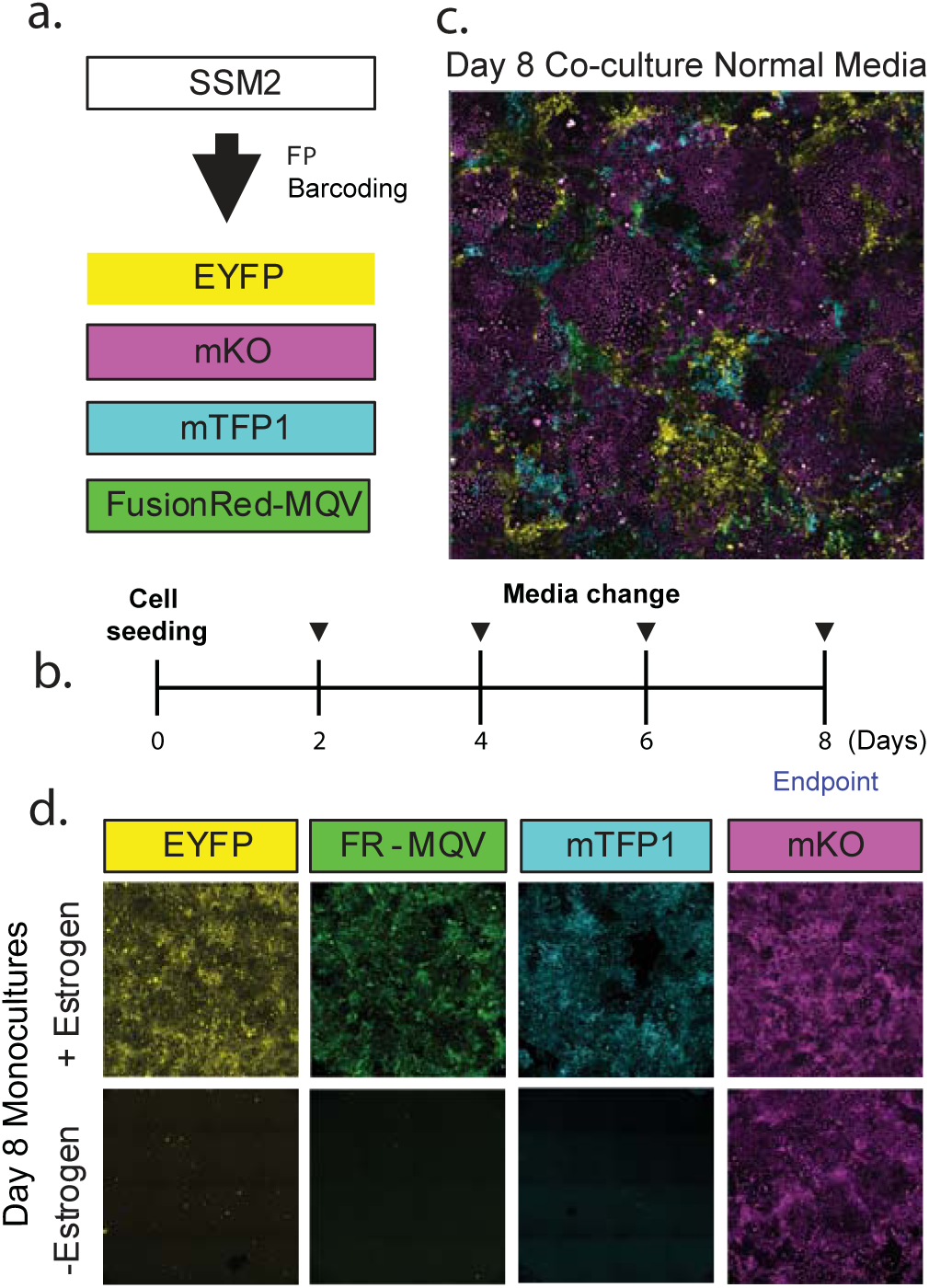
Visualizing clonal dynamics of SSM2 estrogen independence. **(a)** Four different spectrally resolvable SSM2-FluorescentProtein (FP) labeled cell lines were generated (SSM2-eYFP, SSM2-mKO, SSM2-mTFP1, SSM2-FusionRed-MQV) using CMV-FP-PiggyBac transposon integration and FACS purification. **(b)** Timeline of SSM2-FP proliferation assay. Endpoint of day 8 determined by confluency in normal media conditions. **(c)** Representative image of SSM2-FP co-culture at day 8. Cells reach confluence in normal media and FPs can be reliably deconvoluted. **(d)** Representative images of day 8 monocultures for each SSM2-FP cell line. Top row images are cells grown in normal media and bottom row images are cells grown in estrogen-depleted media. All SSM2-FP cell lines grow to confluence in normal media, but only the SSM2-mKO cell line grows to confluence in estrogen depleted media.

## DISCUSSION

Breast cancer is the second most common cause of cancer-related death among women in the US (26). ER^+^ carcinomas account for 79% of all incidences and 67% of all breast cancer mortality (27). However, approximately 30% of ER^+^ breast cancers do not respond to anti-estrogen hormone therapies(28). While the animal models can be used to decipher the mechanisms of anti-estrogen drug resistance, only a few mouse models have been reported to phenocopy of ER+ breast cancer(16, 20, 21, 28). Furthermore, only a few groups have successfully developed experimental models that exploit an ovarian hormone- and estrogen-dependency of murine mammary tumor growth(23, 29, 30).

The experiments described here were designed to determine how one type of ER^+^ positive luminal type mammary tumor becomes estrogen independent tumors. We previously used the genetically modified *Stat1^-/-^* mouse model which spontaneously develops ER^+^ luminal type mammary tumors(20, 21). The mammary tumor developed in this *Stat1^-/-^* mouse initially regressed but later resumed growth following ovariectomy treatment (23). Additionally, syngeneically and orthotopically transplanted STAT1*-null* primary tumor or *Stat1^-/-^* mouse-derived mammary tumor cell line SSM2 failed to grow in ovariectomized *Stat1^-/-^* mouse(20). In this study, we attempted to syngeneically and orthotopically transplant STAT1-null mammary tumors and SSM2 cells into 129S6/SvEv wild-type mice to model an anti-estrogen drug resistance. Interestingly, both grown transplanted SSM2 and STAT1-null primary tumors continued to grow after ovariectomy treatment in our study. Given the continued ovary-independent growth, we explored the possible mechanisms. Our analysis of SSM2-derived transplanted tumors revealed a phenotypic shift from a luminal to a basaloid subtype following ovariectomy treatment, suggesting two key implications. First, the SSM2 transplant model can serve as a phenocopy of luminal-type breast cancer. Second, SSM2 transplantation combined with ovariectomy can also be utilized as a model for anti-hormonal therapy-resistant breast cancer.

The SSM2-derived transplanted tumor initially exhibits an ER+ luminal phenotype. However, two weeks after ovariectomy, small clusters of CK14-positive metaplastic basaloid cells emerge within the tumor. These basaloid tumors display high proliferative activity and contain high percentages of ER- and p53-positive cells, which form irregular clusters frequently composed of cells with elongated and ellipsoid nuclei. By eight weeks post-ovariectomy, approximately 90% of the tumor cells have transitioned into a metaplastic basaloid phenotype. These tumors represent the ultimate form of mammary metaplasia and epithelial-mesenchymal transition (EMT), processes known to be associated with poor prognosis in mammary tumors (12). High p53 expression levels is known to strongly link to EMT (31), whereas p53 loss of function has been shown to induce high aromatase levels (32). Studies on human breast cancer specimens have demonstrated a significant association between p53 accumulation and resistance to aromatase inhibitors in ER+ breast cancer (33, 34). Our findings reveal that EMT-like metaplastic tumors are correlated with a high percentage of p53+ tumor cells in both SSM2 and primary tumor transplants, particularly under ovariectomy conditions. This experimental condition closely parallels the state of ER+ breast cancer in patients receiving aromatase inhibitor therapy (33, 34).

The phenotypic shift from luminal to basaloid tumors appears to reflect selection of a minor subclone present in the SSM2 cell line. Initially, p53 expression is present in a low percentage consistent with a p53 “wild type” expression pattern. Sequencing of the SSM2 cells and the luminal phenotype tumors revealed a p53 S12R variant, predicted to be non-deleterious. Following ovariectomy, as the tumor transitions into a basaloid phenotype, p53 is stabilized resulting in an increased tumor cell IHC percent positive (greater than 60%) consistent with a p53 functional mutation. Sequencing shows a new p53-C132W variant. Notably, p53 C132W mutation corresponds to a common “hot spot” p53 mutation found human tumors. Similarly, mutations in the prolactin receptor (Prlr) follow a distinct pattern. The p.R319Lfs*6 mutation is present in both untreated SSM2 tumors and p53⁻ˡᵒʷ SSM2 tumors post-ovariectomy, suggesting that this mutation is an intrinsic feature of SSM2 cells. In contrast, the p.E318Sfs*3 mutation is detected only in p53-C132W SSM2 tumors post-ovariectomy, implying that it emerges under conditions of ovarian hormone (estrogen) deprivation. These findings suggest that a minor cell population within SSM2 tumors, carrying the p53 p.C132W and/or *Prlr* p.E318Sfs*3 mutations, become highly proliferative in the absence of ovarian hormones, driving the transition to a basaloid phenotype. This hypothesis is further supported by the survival of fluorescence protein-barcoded SSM2 cells in estrogen-depleted culture media. Although further validation is needed to confirm whether the same gene mutation patterns occur in cultured SSM2 cells, these findings strongly suggest the presence of a minor subpopulation resistant to estrogen deprivation.

Aside from p53 and Prlr mutations, gene mutation analysis reveals distinct patterns in the SSM2-derived transplanted tumor. These mutations can be categorized into three major groups: (1) ovariectomy treatment-dependent mutations (Fig. 4: Ca2 and Cb1), (2) luminal p53⁻ˡᵒʷ tumors (Fig. 4: Ca3 and Cb2), and (3) basaloid tumors (Fig. 4: Ca5, Cb3, and Cb4). The mutation patterns suggest that SSM2 cells comprise multiple subpopulations. Ca2 and Cb1 mutations were exclusively present in SSM2 tumors with ovariectomy (Ca2) and without ovariectomy (Cb1). This distinct pattern supports the existence of ovarian hormone-sensitive and - insensitive populations, with the insensitive population demonstrating increased proliferation or survived in the absence of ovarian hormones. In luminal p53⁻ˡᵒʷ tumors (Ca3 and Cb2), the mutation patterns indicate that luminal-type SSM2 cells persisted under ovariectomy treatment in tumors with Ca3 mutations, while a subpopulation of SSM2 cells gave rise to tumors harboring Cb2 mutations. Notably, *Esr1* p.G78A and p.K184R mutations were identified in Cb2. These mutations are in the AF1 and DBD domains, respectively, both of which are known to regulate ER’s transcriptional activity (35, 36). Since these ER mutations were found in tumors under ovariectomized conditions, the mutant ER proteins may exhibit activity independent of estrogen binding. Conversely, most basaloid tumors likely arose from a rare pre-existing subpopulation that expanded after ovariectomy treatment. This is supported by the observation that only one mutation was retained in Ca5, whereas 23 unique mutations were identified in basaloid-specific tumors (Cb3 and Cb4). Although the direct relationship between these mutations and the luminal-to-basal transition remains to be elucidated, we anticipate that further investigation will clarify the connection between gene mutation patterns, subpopulations, and their roles in tumor progression in the near future.

A recent comprehensive genomic landscape study analyzing 22 human primary ER+ breast tumor specimens before and after aromatase inhibitor therapy has highlighted clonal complexity, with the emergence of newly detected or enriched independent tumor clones post-treatment (35). The study stated that “*precision medicine approaches based on genomic analysis of a single specimen are likely insufficient to capture all clinically significant information*” (35). Our findings support this concept, demonstrating that independent clonal selection of metaplastic cells can occur following ovarian hormone loss. Here, we report that deleterious *Trp53* mutations, along with an increased percentage of p53-positive cells, are associated with ovarian-independent tumor cell population, which differs from the major cell populations in the initial tumors. Our analysis identified small clusters of metaplastic cells within the dying luminal tumor epithelium, suggesting that these clusters may serve as the precursor cells for ovarian-independent tumors, which subsequently persist and expand in long-term models. Therefore, our findings reinforce the notion that current approaches, including initial genomic and pathological analyses, may be insufficient to predict whether ER+ breast cancers will ultimately develop hormone-independent tumors. If true, we may be able to detect minor basaloid and/or mutant p53 subclones in some ER+ breast cancers that predict resistance to anti-hormonal therapies. For example, increased sequence depth may reveal p53 mutations in ER+ breast cancers present at low/very low percentage/allele frequency in the higher risk cancers.

## ACKNOWLEDGEMENT

Research in this study was supported by National Health Institute U01CA196406 (A.D.B), K01OD030518 (H.M.), and UCD Comprehensive Cancer Center Support grant under NIH P30CA093373 (H.M., A.D.B.).

## MATERIALS AND METHODS

### Animals

129S6/SvEvTac-*Stat1^tm1Rds^* (Tm(*Stat1^-/-^*)) mice (20) were contributed by the Schreiber lab, and wild-type 129S6/SvEv (129SvEv WT) mice were purchased from Taconic Farms. Mice were housed in a vivarium under NIH guidelines and all animal experiments followed protocols approved by the UC Davis Institutional Animal Care and Use Committee (IACUC).

### Orthotopic syngeneic transplantation

Primary tumors from *Stat1^-/-^* mice were removed surgically. When mammary tumors developed from donor *Stat1^-/-^*mice, 1-2 mm^3^ in size of tumors were dissected from the number 2 mammary fat pad. Collected STAT1-null primary tumors were processed for histopathology as well as transplanted into the uncleared number 4 and 9 fat pads of 4-week-old 129SvEv WT recipient mice. All surgical orthotopic transplantation was performed under Nembutal anesthesia (60 mg/kg) followed by post-surgical analgesia (Buprinex; 0.05 mg/kg). All mice were euthanized using an overdose of Nembutal (120 mg/kg) prior to tumor collection and fixation of tissues.

SSM2^UCD^ cells were cultured in DMEM/F12 containing 10% FBS, 1% L-glutamine, 1% penicillin-streptomycin, 50 μM 2-mercaptoethanol, 0.3 μM hydrocortisone, 5 μg/mL insulin, and 10 ng/mL transferrin. Similar to *Stat1^-/-^* primary tumor transplantation, for orthotopic syngeneic SSM2 transplantation *in vivo*, 1 x 10^5^ SSM2 cells were transplanted into the uncleared number 4 and 9 fat pads of 4-week-old 129 SvEv wild-type recipient mice under anesthesia. To calculate tumor growth *in vivo*, tumor volumes were measured using a caliper with the equation, (L x W^2^)/2. To remove the tumors, mice were first anesthetized using Nembutal (60 mg/kg) and tumors removed from live, anesthetized animals. Following removal, the mice were euthanized using an overdose of Nembutal (120 mg/kg). All tissues were immediately cut into 1-2 mm slices using a razor blade or scalpel and placed immediately into fixative for further histopathology exam.

### SSM2 cell line generation

SSM2 cells were maintained in DMEM media (High glucose, glutamine, phenol red) with 10% FBS and 1% anti/anti (Thermo Fisher) at 37°C in a humidified incubator with an atmosphere of 5% CO_2_. For stable expression of FPs, cells were transfected with CMV-FP-PiggyBac transposon and hyPBase transposas (gift from Christopher Newgard) plasmids using Lipofectamine 3000 reagent. Cells were allowed to recover for 3-7 days and then selected for FP expression using FACS. Cell purity was defined as >90% of cells in expressing the FP. Cell lines underwent serial FACS as needed until purity threshold obtained. CMV-FP-PiggyBac transposons were generated by cloning IDT synthesized FP fragments into the p-MVP gateway cloning system (Addgene #1000000155) as described previously (36).

### SSM2-Proliferation Assay

Estrogen-independence of each SSM2-cell line was defined by the ability of cells to grow in estrogen-depleted media (10% charcoal-stripped FBS in phenol red-free DMEM with 1% anti/anti). The proliferation assay was performed in a 48-well glass-bottom plate (MatTek). SSM2-FP cell lines were seeded into normal or estrogen-depleted media in monoculture or equal concentration co-cultures. Each condition was plated in triplicates. Media was changed every other day, and cells were monitored for confluency. Endpoint was reached at day 8, at which point the plate was imaged pre-fixation. Imaging was performed using a Zeiss LSM 880 (Carl Zeiss, Inc., Thornwood, NY, USA) laser scanning confocal microscope equipped with an incubated chamber to maintain optimal conditions for live-cell imaging. Prior to imaging live cells, the incubator’s temperature was set to 37°C, humidity control was activated, and the CO_2_ concentration was adjusted to 5%. All conditions were imaged during the same imaging session.

### Histopathology and immunochemistry

Tissues were fixed in 10% neutral buffered formalin at room temperature for 24 h then placed in 70% ethanol until processed, which was normally within 24 h. A Tissue-Tek VIP autoprocessor (Sakura, Torrance, CA) was used to process tissues that were then embedded in Paraplast (melting temperature 56–60°C), sectioned to 4 μm and mounted on glass slides. Sections were stained with hematoxylin and eosin (H&E) for pathologic analysis. Sections were also stained for estrogen receptor (ER) and p53 immunohistochemistry (IHC). Primary antibodies used in the study were rabbit polyclonal anti-ER (1:100; SC-542) and rabbit polyclonal anti-p53 (1:1000; SC-6243) from Santa Cruz Biotechnology, Inc. (Santa Cruz, CA). For ER staining, mouse uterine tissue was used as a positive control and for p53, mouse (Pb - SV40 Lg/Sm Tag) prostate with papillary hyperplasia was used. Antibody deletion controls were used to confirm specific staining. Antigen retrieval was performed on tissue sections using a Decloaking Chamber (Biocare Medical, Concord, CA) with citrate buffer at pH 6.0, 125°C and pressure to 15 psi. The total time slides were in the chamber was 45 min. Incubations with primary antibodies were performed at room temperature overnight in a humidified chamber. Normal goat serum was used for blocking, biotinylated goat anti-rabbit (1:1000) was the secondary antibody used with a Vectastain ABC Kit Elite and a Peroxidase Substrate Kit DAB (Vector Labs, Burlingame, CA) used for amplification and visualization of signal, respectively.

### Sequencing

Total DNA was purified from FFPE tissues for sequence analysis via BGI Genomics. Adapter sequence was trimmed with cutadapt. Reads were aligned to mm10 mouse reference using BWA, and PCR duplicate reads were marked with sambaba. Base quality recalibration was performed with Genome Analysis Took Kit 4. Mutation was called with Mutect2. Annotation was performed with VEP. Germline variant was filtered using cross sample identification and variant allele frequency. For Sanger sequencing, Total DNA was also purified from FFPE tissues. Total DNA was extracted and amplified using Sequalprep long PCR kit (Life Technologies). Purificants were submitted to UCD sequencing facility and then analyzed using Genecodes Sequencher software. All potential genetic mutations were re-PCR amplified, purified, and sequenced to confirm the mutant candidate was not a PCR induced artifact.

### Image analysis

All slides were scanned to produce digital whole slide images (WSI) using either an Aperio ScanScope XT or AT2 and stored in the Aperio Spectrum database customized for laboratory workflow. Quantitative image analysis was performed using Aperio Imagescope software, based on the FDA approved algorithms supplied by the manufacturer. For quantitative image analysis of each IHC, regions of interest on WSI were validated by two pathologists (ADB, RDC) and analyzed using the Aperio IHC Nuclear algorithm. Percent positive nuclei for p53 was calculated automatically by the algorithm.

#### Multiplex Immunohistochemistry and its imaging analysis

Formalin-fixed, paraffin-embedded (FFPE) tissue sections, prepared at a thickness of 4 microns, were stained for a multiplex immunohistochemistry using the Leica BondRX autostainer (Leica Biosystems). The primary antibodies used in this study included anti-p53 (Bioss), anti-ER (Santa Cruz Biotechnology and Millipore), anti-Krt14 (CK14; ABclonal), and anti-Ki67 (Cell Signaling Technology). For secondary detection, 1x Opal Anti-Ms+Rb HRP (Akoya Biosciences) was used, and signal amplification was achieved with TSA-conjugated Opal dyes (Akoya Biosciences), including Opal-690 for p53, Opal-620 for ER, Opal-570 for CK14, and Opal-520 for Ki67. The antibody-Opal dye pairings, concentrations, and the sequential order of staining were optimized based on previously established protocols (37) to maximize signal-to-noise ratios and achieve balanced signal intensities. Stained tissue sections were scanned using the Vectra/Polaris multispectral imaging system (Akoya Biosciences), and the resulting im3 files were converted to multilayered TIFF files using inForm software (Akoya Biosciences). Image analysis was performed in QuPath software (38) using its cell detection feature to segment cells and quantify marker expression levels in the nucleus (p53, ER, and Ki67) and cytoplasm (CK14). Data analysis and pie chart generation were conducted using Python with the pandas and matplotlib.pyplot libraries, while bar graphs and statistical analyses were performed using GraphPad Prism.

---

**Supplementary Figure 1.**
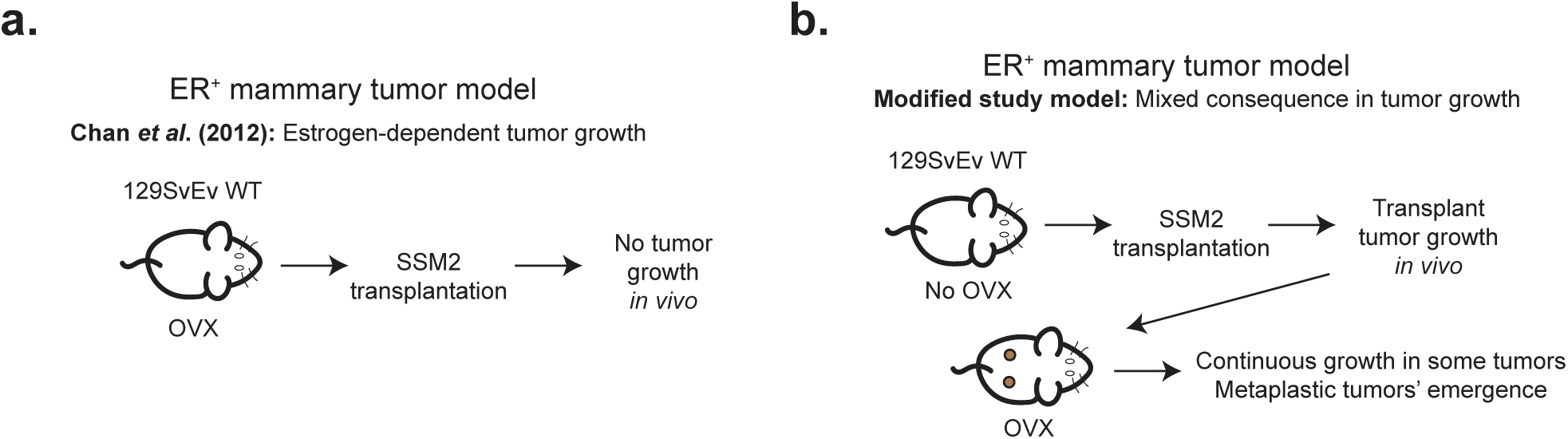
Procedures for generating SSM2 cell transplants as an ER+ mammary tumor model. (**a**) A previously established procedure for using SSM2 cell transplants as a model for ER+ mammary tumors, as reported by Chan et al. (2012), demonstrated that SSM2 cells failed to form tumors when transplanted into ovariectomized mice. (**b**) In the current study, we developed a modified model by syngeneically orthotopically transplanting SSM2 cells into the mammary fat pad of mice without prior ovariectomy. Tumor growth was monitored, and once tumors were established, the mice were ovariectomized to study the fate and phenotypic changes of the SSM2 tumors.

**Supplementary Figure 2.**
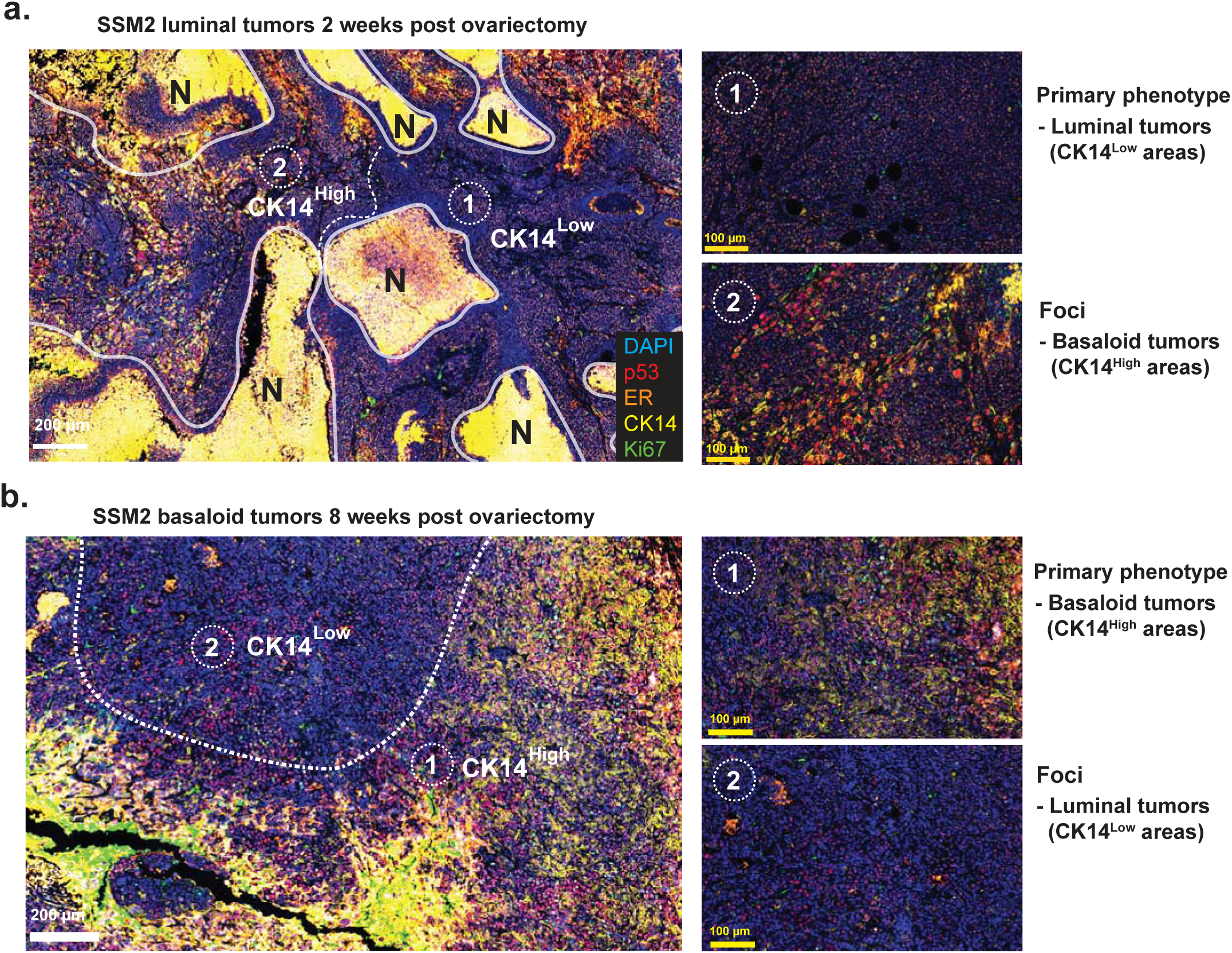
Multiplex IHC images of tissues from (**a**) SSM2 luminal tumors at two weeks post-ovariectomy and (**b**) SSM2 basaloid tumors at eight weeks post-ovariectomy. For each tumor, the (1) primary phenotype and (2) foci are highlighted. Markers are visualized as follows: p53 (red), ER (orange), CK14 (yellow), Ki67 (green), and DAPI (blue). Areas of necrosis are indicated by “N.”

**Supplementary Figure 3.**
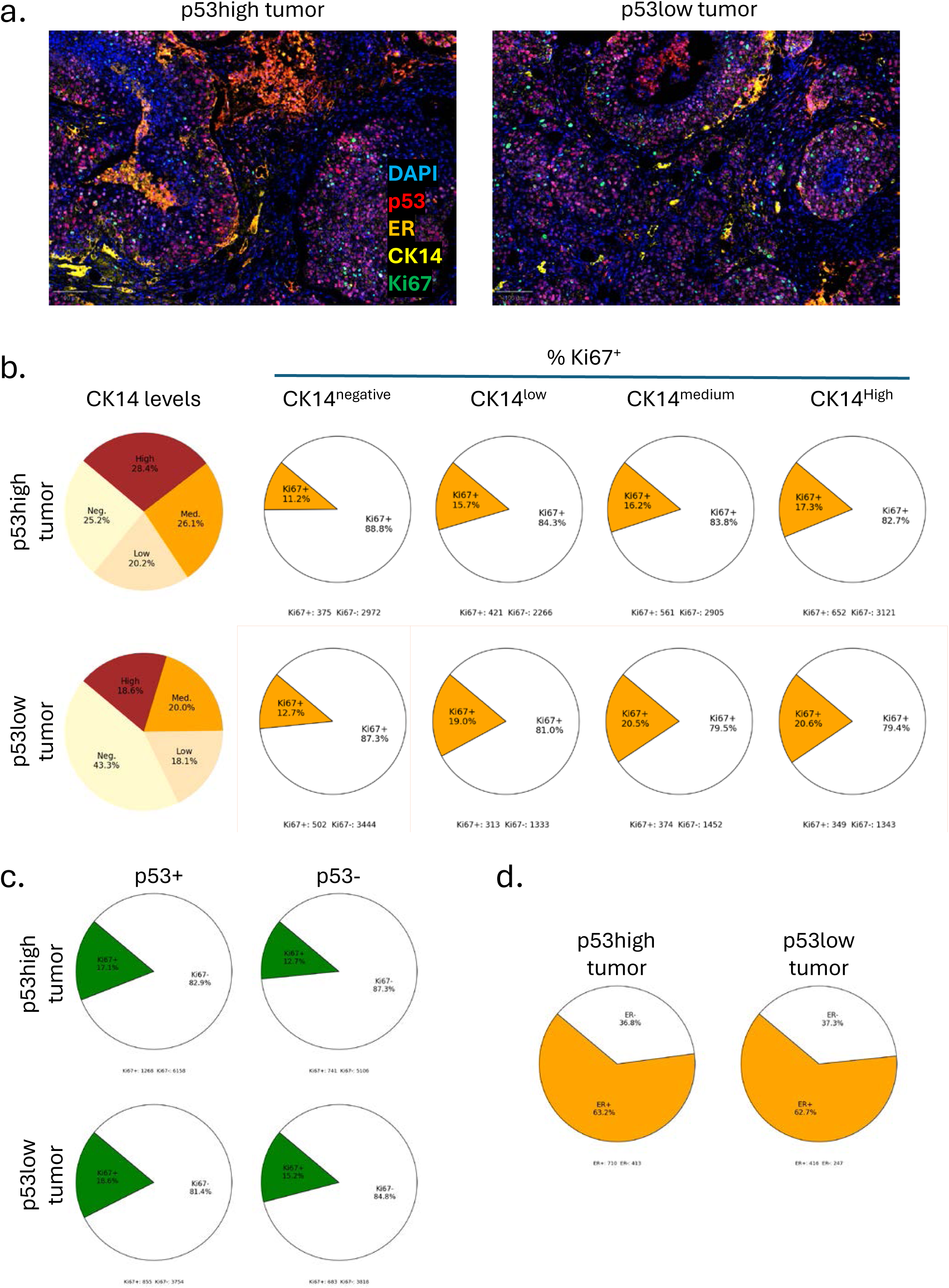
Multiplex IHC analysis of p53-High and p53-Low primary tumors grown in the mammary fat pad of ovariectomized mice. Tissue sections of p53-High and p53-Low primary tumors grown in the mammary fat pad of ovariectomized mice were stained using a multiplex IHC panel to detect p53, ER, CK14, and Ki67. (**a**) Representative multiplex IHC images of p53-High and p53-Low primary tumors. The markers are visualized as follows: DAPI (blue), p53 (red), ER (orange), CK14 (yellow), and Ki67 (green). Scale bar = 100 µm. (**b**) Pie charts showing the percentages of CK14 expression levels (CK14-negative, CK14-Low, CK14-Medium, and CK14-High) in p53-High and p53-Low tumors. Each pie chart also displays the percentages of Ki67-positive (orange) and Ki67-negative (white) cells within each CK14 expression level category. (**c**) Cell proliferation status of p53-positive and p53-negative cells in p53-High and p53-Low tumors. Pie charts depict the proportions of Ki67-positive (green) and Ki67-negative (white) cells. (d) Pie charts illustrate the percentages of ER-positive (orange) and ER-negative (white) cells in p53-High and p53-Low tumors. Cell counts are displayed below each pie chart.

**Supplementary Table 1.**
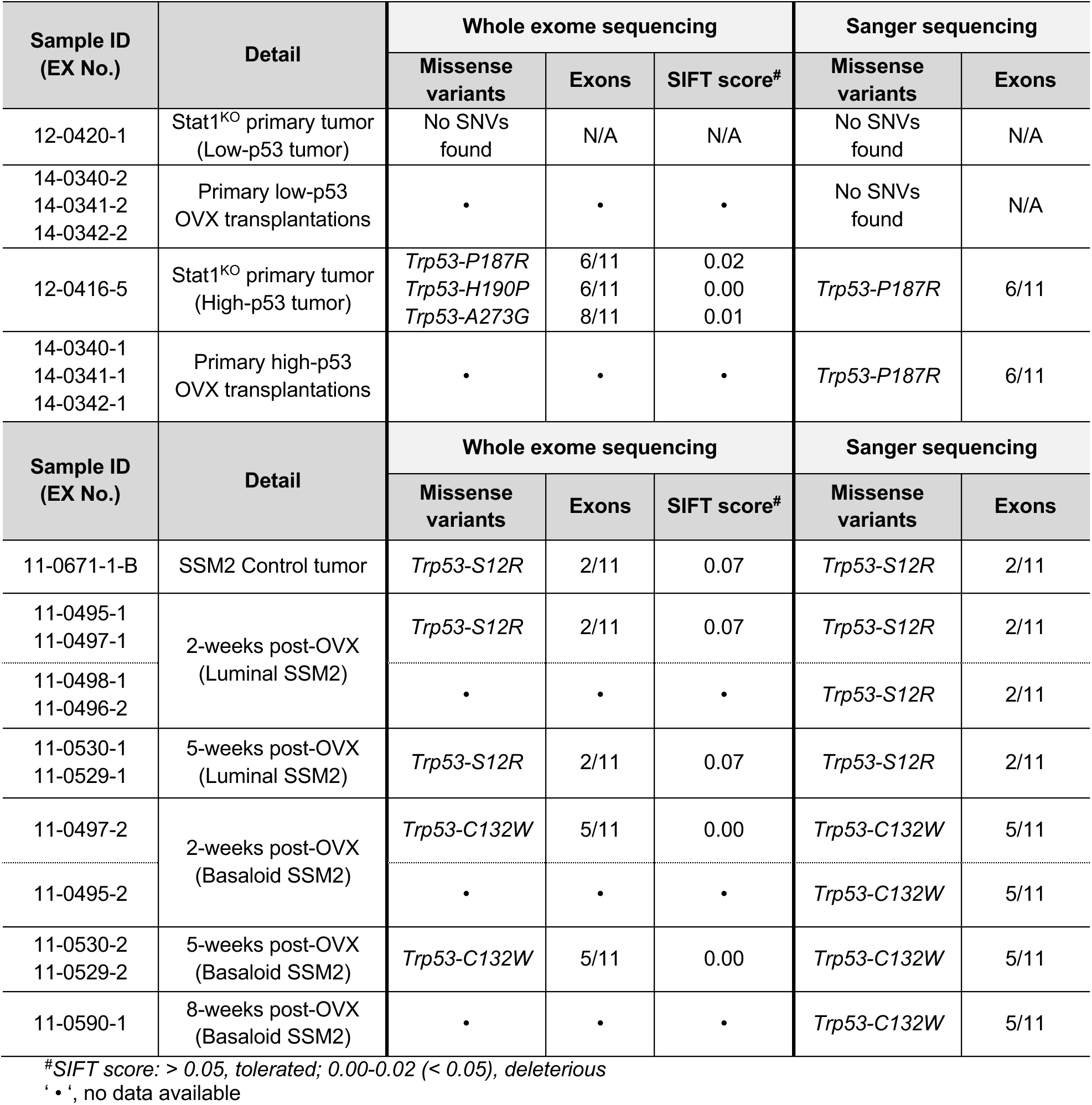
Summary of *Trp53* mutations in STAT1-null primary and transplanted tumors, and transplanted SSM2^UCD^ tumors before and after ovariectomy.

**Supplementary Table 2.**
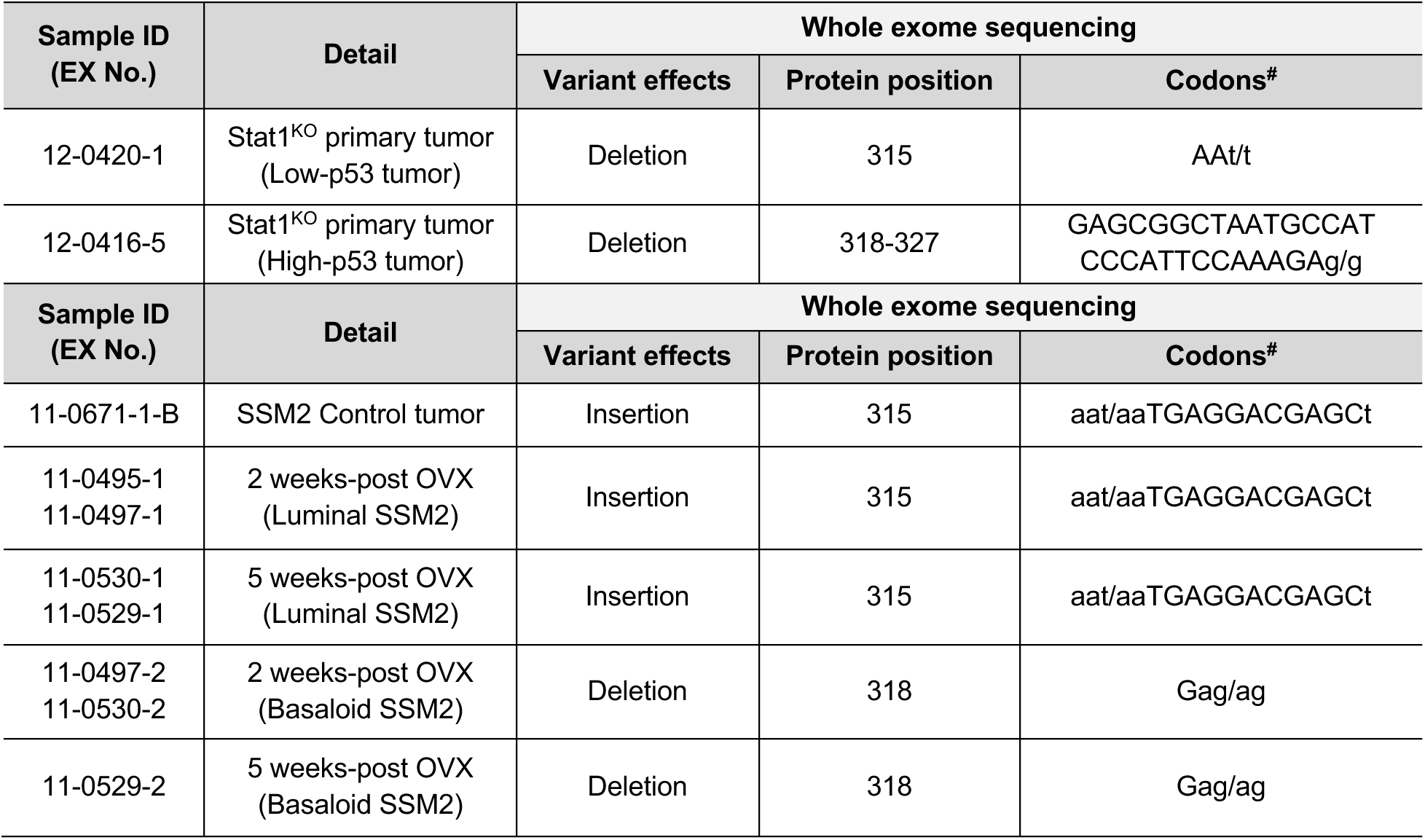
Summary of *Prlr* mutations in STAT1-null primary and transplanted tumors, and transplanted SSM2^UCD^ tumors before and after ovariectomy.

